# Fitness impacts of *Plasmodium* vary by host age and sex in a North American songbird

**DOI:** 10.1101/2025.10.27.683448

**Authors:** Katherine M. Talbott, Oxana V. Gorbatenko, Cierra McKoy, Ellen D. Ketterson

## Abstract

Birds infected with the malaria-causing parasite *Plasmodium* comprise a popular model in evolutionary ecology, yet *Plasmodium’s* impact on host survival and reproduction remains unclear in wild bird populations where *Plasmodium* is historically endemic. Recent research in endemic host populations shows that *Plasmodium* reduces survival in juveniles, but not adults. Additional research is needed to understand 1) age-based variation in the impact of *Plasmodium* on host reproduction, and 2) sex-based variation in the impact of *Plasmodium* on survival and reproduction. To this end, we leveraged a long-term dataset from a dark-eyed junco (*Junco hyemalis carolinensis*) population with a high prevalence of *Plasmodium*. Juvenile males had lower probability of *Plasmodium* infection than juvenile females, indicating either elevated resistance or mortality in young males before the juvenile stage. Second year females had a lower probability of infection than females of other age classes, which suggests elevated mortality in infected yearling females after their first breeding season. Model comparison did not identify a direct relationship between *Plasmodium* infection and adult return (a proxy of survival), yet surprisingly, females paired with infected males in the previous breeding season had a higher probability of return. There was no relationship between *Plasmodium* infection and juvenile return, but returners with relatively higher parasite loads as juveniles had shorter lifespans than those without juvenile infections. Among adults, *Plasmodium* infection did not predict fledging success or fledgling number. However, returning juncos with relatively heavier bodies and higher parasite loads as juveniles had an increased probability of breeding in their first adult year. Combined with the shorter lifespans observed in infected juveniles, this suggests a terminal investment strategy for juncos contracting *Plasmodium* early in life. Our results indicate that the impact of *Plasmodium* on host fitness likely varies by host age at first infection and may exacerbate physiologically stressful life stages.

## Introduction

Birds infected with the malaria-causing parasite *Plasmodium* comprise a popular model in evolutionary ecology [1,2], yet knowledge of the extent to which this parasite influences differential survival and reproduction of its avian hosts is incomplete. Songbird mortality events are well documented in *Plasmodium*-naïve populations following the introduction of *Plasmodium*-vectoring mosquitoes [3–6], but recent studies suggest *Plasmodium* may also impact fitness in host populations with a long coevolutionary history with the parasite. For example, chronic *Plasmodium* infections have been shown to reduce longevity [7–11] and reproductive output in adults of some songbird species [7,11–16]. In contrast, the strongest selective pressure of this parasite is predicted to occur through initial infections in juvenile songbirds [6,17], yet relatively little is known about these impacts. After hatching naïve to *Plasmodium*, a young bird’s first infection necessarily develops into the acute stage, which is associated with higher parasite loads and more severe health impacts compared to chronic infection [18]. Empirical studies testing the hypothesis of age-based variation in selective effects of *Plasmodium* in wild birds are few, likely because of the logistical challenges of capturing young, infected songbirds in sufficient numbers for study. One such recent study in house sparrows (*Passer domesticus*) showed that while high-intensity infections of *P. relictum* were negatively associated with survival among sparrows of all ages, such infections in juveniles were predictive of population decline, while high-intensity infections of adults were not [19].

Building upon this, additional research is needed to determine whether these age differences in host survival attributable to *Plasmodium* also vary by sex, and whether they extend to host reproductive success.

With respect to sex, there are several reasons why adult females might be expected to show larger *Plasmodium*-related reductions in survival compared to males. Mark-recapture studies in temperate passerine populations have shown higher annual survival in adult males, compared to females [20–22]. This sex bias in longevity has been attributed to relatively higher reproductive effort and vulnerability to predators during incubation in females [22,23], as well as the expression of deleterious alleles in females as the heterogametic sex [21]. In addition, females may experience trade-offs between immune and reproductive investment during the breeding season [24–27], potentially leaving them more vulnerable to the impacts of *Plasmodium*. Therefore, we predicted that adult females would show larger *Plasmodium*-related reductions in survival than adult males. On the other hand, we might expect the opposite pattern among younger birds, with greater *Plasmodium*-related mortality among males. For example, male nestlings are sometimes more heavily impacted by environmental factors such as parasitism [28,29], poor nest site quality [30], and enlarged broods [31,32], compared to females. Therefore, we predicted larger *Plasmodium*-related reductions in survival in juvenile males, compared to juvenile females.

We might also expect age-based variation in the impact of *Plasmodium* on host reproductive success. While such impacts are well-studied in adult birds, relatively few have asked whether parasitism during the juvenile stage influences future reproductive success [33]. For example, the metabolic cost of a juvenile’s immune response to acute infection could reduce nutrient and energy reserves [17,18,28,34], which are critical resources for its first breeding season. In short-lived passerines, foregoing or delaying first year breeding could have considerable impact on individual lifetime reproductive success; thus, the selective pressure of *Plasmodium* on host reproduction could be stronger in juveniles than in adults. Hence, we predicted that *Plasmodium*-related decreases in host reproduction may be associated with juvenile infections, even if they are not evident in infected adults.

To test these predictions, we investigated demographic variation in the relationship between *Plasmodium* parasitism and host fitness using data from a long-term ecological study of dark-eyed juncos (*Junco hyemalis carolinensis*, hereafter ‘junco’). First, we explored demographic patterns in the probability of *Plasmodium* infection and in parasite load. Then, we used a model comparison approach to investigate the ability of *Plasmodium* infection status and load to predict metrics of survival and reproduction in both adult and juvenile juncos. With this approach, we specifically tested the hypotheses that 1) *Plasmodium* parasitism reduces the fledging success (i.e., ability to raise at least one young to the point of leaving the nest) and fledgling number in breeding adults, and 2) *Plasmodium* parasitism in juvenile hosts reduces the probability of first-year breeding attempts. We also used recapture (‘return’) in years following sampling as a proxy of survival to test the hypotheses that 3) *Plasmodium* parasitism reduces survival in adults and 4) both survival and lifespan in juveniles. Better understanding demographic variation in fitness impacts of *Plasmodium* will help refine our predictive framework for investigating songbird-*Plasmodium* coevolution and inform conservation priorities for songbird populations threatened by *Plasmodium*.

## Materials and Methods

### Field sampling

This study focuses on a non-migratory population of dark-eyed juncos (*Junco hyemalis carolinensis*) near Mountain Lake Biological Station (MLBS) in Giles County, VA, USA, with a high prevalence of *Plasmodium* (Supporting Text S1 Section A). Morphometric data and junco blood samples were collected during annual censuses as part of a longitudinal study on junco evolutionary ecology [35]. An early breeding season adult census takes place from mid-April through mid-May, and a late breeding season census of independent young, termed ‘juveniles’, occurs from July through early August (Supporting Text S1 Section B). Following the adult census, researchers actively searched for nests on the study site and identified social mates by observing interactions at nest sites, noting unique band combinations. Fledging success was defined as an adult pair raising one or more offspring to the point of leaving the nest within a breeding season. Pairs with two or more documented nest failures and no known fledglings during a specific breeding season were scored as ‘unsuccessful’; only pairs fitting the criteria of ‘successful’ or ‘unsuccessful’ were included in our analyses. Furthermore, only one sample per bird was used for analyses; that is, there were no instances of repeated sampling across years for any individual (Supporting Text S1 Section C). Long-term data from marked individuals was used to infer junco age, survival, and lifespan. Adult age category was divided into yearlings (i.e., adults in their first breeding season, which hatched the prior year), two-year-olds, and birds three years of age or older. Recognizing that failure to return may be due to mortality or dispersal, we used return as our best available and conservative proxy of overwinter survival. Because dispersal is generally female-biased in juncos [36,37], we ran analyses of junco return separately by sex.

### Plasmodium diagnostics and parasite load quantification

We used quantitative polymerase chain reaction (qPCR) to detect *Plasmodium* infections in junco blood samples and to assess relative amounts of parasite DNA and host DNA, scaled to the bird with the highest proportion of parasite DNA (Supporting Text S1 Section D). Parasite load was quantified by multiplying relative quantification values by 100,000 and then rounded to the nearest integer to facilitate use in generalized linear models (see ‘Statistical analysis’ below). To reduce the impact of outliers on model residuals, juvenile and adult loads were Winsorized at 90%; that is, values above the 90^th^ percentile were assigned the 90^th^ percentile value [38]. Because of the phenology of *Plasmodium*-vectoring mosquitoes in the study location, we assume that adult juncos testing positive for *Plasmodium* are in a chronic state of infection, while juveniles testing positive for *Plasmodium* are likely in the acute stage of their first *Plasmodium* infection (Supporting Text S1 Section A).

### Statistical analysis

All statistical analyses were conducted in R (version 4.3.1). We used a generalized linear model (GLM) with a binomial family and logit link to ask whether the interaction of sex (male or female) and bird age class (adult or juvenile) predicted the probability of *Plasmodium* infection. We used a GLM with a negative binomial family and log link to ask whether the interaction of sex and age class predicted *Plasmodium* load in *Plasmodium*-infected birds.

Next, we used a model comparison approach to explore ecological predictors of *Plasmodium* infection status and load, with separate analyses for juvenile and adult juncos. Finally, we used a similar model comparison approach to identify predictors of adult junco fitness metrics (fledging success, number of fledglings, and return to the MLBS population) and juvenile junco fitness metrics (return to the MLBS population, lifespan, and first year breeding). The family (negative binomial, binomial, or Poisson) for each series of GLMs was selected based on the distribution of the dependent variable and best model fit, and canonical link functions were used in all cases.

For each model comparison, the model list included one univariate model per predictor along with multivariate models containing a combination of terms. Predictors included those related to *Plasmodium* infection (i.e., *Plasmodium* infection status, *Plasmodium* load, or a mate’s *Plasmodium* infection status or load), junco ecology or body size (e.g., body condition, wing length, age category, adult fledging success), or environmental conditions (i.e., year). This approach provides an ecologically relevant context for interpreting the importance of *Plasmodium* as a predictor of songbird fitness and helps disentangle interactions between *Plasmodium* parasitism and ecological factors such as sampling year, host sex, or host age.

To reduce likelihood of overfitting, all candidate models were limited to no more than one predictor per 10 observations. For each analysis, we compared candidate models by using the Akaike Information Criterion (AICc) for small sample sizes [39], using the package *bbmle* [40]. We considered models within two ΔAIC to be competitive, and these models comprised top model sets. For each top model set, we confirmed normality of model residuals by inspecting Q-Q plots with the R package *DHARMa* [41] and confirmed the absence of collinearity using the *Performance* package [42]. For analyses in which the model with the lowest ΔAIC was also the most parsimonious (i.e., had fewest predictors), we considered this to be the best-performing model (‘top model’). For analyses with multiple models within two ΔAIC, we removed models with simpler, higher-ranking (by weight) variants from the top model set [43] and then used conditional averaging on the remaining top models to produce estimates for each variable. We elected to use conditional model averaging over full model averaging to allow detection of subtle relationships [44]. For estimates that did not span zero, we included odds ratios. See S1 Supporting Tables for details of all competitive models for each analysis.

## Results

### A. Which factors predict Plasmodium infection status and load?

#### Among all juncos, does *Plasmodium* infection and/or load vary with age class or sex?

Among all juncos, age class (i.e., adult [n = 104] or juvenile [n = 145]) was a significant predictor of the probability of *Plasmodium* infection (X^2^ = 9.68, df = 1, p = 0.002). Odds of infection were 2.42 times higher in adults compared to juveniles (estimate = 0.88 ± 0.39 SE; see Figure 1A). However, infection status did not vary by sex (X^2^ = 0.98, df = 1, p = 0.32), nor was there an interaction between sex and age class (X^2^ = 0.02, df = 1, p = 0.90).

**Figure 1.** Sex-and age-based variation in *Plasmodium* parasitism in dark-eyed juncos (*Junco hyemalis carolinensis*) sampled near Mountain Lake Biological Station (Pembroke, VA) between 2000 – 2009. Adults were sampled during the early breeding season (12 April – 21 May) and juveniles were sampled during summer (30 June – 9 August). Plot A depicts *Plasmodium* prevalence (± 95% CI, 104 adults and 145 juveniles) and B depicts *Plasmodium* load data in infected individuals (data points are jittered for ease of viewing). See Methods and Materials for *Plasmodium* diagnostic details. Age is given in years, with juveniles shown as zero years of age.

Age class was also a significant predictor of *Plasmodium* load (X^2^ = 26.25, df = 1, p < 0.001), with loads 5.13 times lower in infected adults (n = 47) compared to infected juveniles (n = 38; estimate =-1.64 ± 0.43 SE; see Figure 1B). Load did not vary by sex (X^2^ = 0.31, df = 1, p = 0.58), nor was there an interaction between sex and age class (X^2^ = 0.10, df = 1, p = 0.75). See Figure 1B.

#### Among adults, which factors predict Plasmodium infection status?

Focusing only on adults (n = 104), we investigated year (1999, 2000, 2006, or 2009), age category (i.e., 1, 2, or 3+ yrs of age), sex, and the interaction of age category and sex as predictors of the probability of *Plasmodium* infection. The year 1999 was used as the reference level, as this year had the lowest overall prevalence among adults. We retained three top models <2 ΔAICc; see Table A in the S1 Supporting Tables. Conditional model averaging of the three top models showed that, compared to yearlings, odds of infection were 5.88 times higher in juncos three years old or older (estimate = 1.77 ± 0.76 SE). Odds of infection were 3.29 times higher in females than in males (estimate = 1.19 ± 0.77 SE). However, the relationship between the probability of infection and age depended on sex. In females, unlike males, the probability of infection decreased from one to two years of age (estimate =-1.76 ± 0.99 SE). See Figure 1A.

#### Among adults, which factors predict Plasmodium load?

Potential predictors of *Plasmodium* load in infected adults (n = 47) included wing length (centered by sex), sex, age category, and year, as well as the interactions between sex and wing length, sex and age category, and sex and year. Year was the best predictor of *Plasmodium* load; see Table B in the S1 Supporting Tables. Compared to those sampled in 2000, adults had 3.86 times higher loads in 1999 (estimate = 1.35 ± 0.55 SE), 9.05 times higher loads in 2006 (estimate = 2.20 ± 0.51 SE), and 6.42 times higher loads in 2009 (estimate = 1.86 ± 0.49 SE); see Figures 1B and 2.

**Figure 2.** Year-based variation in parasite load among dark-eyed juncos (*Junco hyemalis carolinensis*) infected with *Plasmodium*. A) Variation in load among infected birds (n = 47), with males represented in red and females in blue. Parasite loads are derived from qPCR analysis of DNA extracted from junco blood sampled during the early breeding season; see Methods and Materials for further details on diagnostics. Points represent data for individual points and are jittered for easier viewing.

#### Among juveniles, which factors predict Plasmodium infection status and load?

Focusing only on juveniles (n = 145), we investigated body condition, sex, hatch year (2000, 2003, 2004, 2006, 2007, 2009, 2010, 2012, or 2014), and the interaction between year and sex as potential predictors of *Plasmodium* infection probability. *Plasmodium* prevalence was lowest among juveniles in 2012; therefore, this year was used as the reference level for models including year as a predictor. We retained two models <2 ΔAICc; see Table C in the S1 Supporting Tables. Conditional model averaging of the two top models showed lower odds of juvenile infection in 2012 compared to all other years sampled, with the most extreme difference in 2003, where odds of infection were 25.76 times higher (estimate = 3.25 ± 1.16). Female juveniles had 2.04 times higher odds of infection compared to male juveniles (estimate = 0.72 ± 0.45 SE); see Figure 3 and Table C in the S1 Supporting Tables. See S1 Supporting Analyses Section A for an additional analysis predicting juvenile infection status from hatch date.

**Figure 3.** *Plasmodium* prevalence by year and host sex in juvenile dark-eyed juncos (*Junco hyemalis*) sampled near Mountain Lake Biological Station (Pembroke, VA). Infection status was determined by molecular analysis of juvenile blood sampled during the summer (30 June – 9 August); see Methods and Materials for additional details. Red lines represent *Plasmodium* prevalence among juvenile male juncos and blue represents females.

Using the same approach, we investigated factors predicting *Plasmodium* load among infected juveniles (n = 38). However, none of our models outcompeted the null to explain variation in load; see Table D in the S1 Supporting Tables.

### B Does Plasmodium parasitism predict fledging success in adults?

Predictors of fledging success among social pairs (n = 52) included: parasite load of each parent, infection category (female infected, male infected, both infected, both uninfected), age of each parent, wing length of each parent, and male body condition. The top model showed higher probability of fledging success in males with longer wings (estimate = 0.35 ± 0.16 SE), with 1.43 times higher odds with each additional mm in length. See Table E in the S1 Supporting Tables. We used the same predictors to investigate variation in fledgling number among successful pairs (n = 26); however, none of our models outcompeted the null model (see Table F in the S1 Supporting Tables).

### C. Does Plasmodium parasitism in juveniles predict the probability of first year breeding?

Among returning juveniles (n = 72), potential predictors of first-year breeding (i.e., being observed in association with a nest) included the following data collected during their juvenile year: *Plasmodium* infection status, parasite load, sex, body condition, wing length (centered by sex), and all possible interactions. See Table G in the S1 Supporting Tables. The top model showed a positive interaction of juvenile parasite loads and heavier body condition in increasing the probability of first year breeding (estimate = 0.02 ± 0.01 SE); see Figure 4.

**Figure 4.** Dark-eyed juncos (*Junco hyemalis carolinensis*) near Mountain Lake Biological Station (Pembroke, VA, n = 72) that had both relatively high *Plasmodium* parasite loads and relatively heavy bodies as juveniles had a higher probability of breeding in their first adult year than those that had relatively low loads and/or relatively light bodies as juveniles. Shown are estimates of a binomial generalized linear model with a logit link predicting the probability of first-year breeding from the interaction of juvenile body condition and *Plasmodium* load; lines represent predictions with loads at the 0, 75^th^, 80^th^, 85^th^, 90^th^, and 95^th^ percentiles. Parasite load data reflect qPCR analysis of DNA extracted from junco blood and are given as the relative measure of host and parasite data, scaled by the proportion of parasite DNA detected in the most heavily infected bird (see Methods and Materials for details).

### D. Does Plasmodium parasitism predict adult return?

We investigated adult return separately by sex (n = 52 per sex), using the following predictors: *Plasmodium* infection status, parasite load, mate infection status, mate parasite load, pair infection category, sampling year, fledging success, age category, mate age category, year, and the interaction between fledging success and infection status. We retained two models <2 ΔAICc for female return; see Table H in the S1 Supporting Tables. Conditional model averaging showed 5.00 times higher odds of return in females that paired with a *Plasmodium*-infected male (estimate = 1.61 ± 0.65 SE); see Figure 5A. In addition, females paired with older males had a higher probability of return; compared to females paired with yearling males, those paired with males two years of age had 7.54 times higher odds of return (estimate = 2.02 ± 1.14), and those paired with males three years of age or older had 12.44 times higher odds of return (estimate = 2.52 ± 1.14).

**Figure 5.** Predictors of adult junco (*Junco hyemalis carolinensis*) return to the Mountain Lake Biological Station population (Pembroke, VA) vary by sex. In females (n = 52), the probability of return is higher in females with *Plasmodium*-infected mates than in females with uninfected mates. In males (n = 52), the probability of return is higher in males that successfully raise at least one offspring to independence (i.e., fledging), than in those that do not fledge any young. Shown are the proportion of returning individuals of each sex, accompanied by 95% confidence intervals.

Our top model for male return showed that males which successfully fledged offspring were 3.52 times more likely to return to the population than unsuccessful males (estimate = 1.26 ± 0.58 SE). See Figure 5B, as well as Table I in the S1 Supporting Tables.

### E. Does Plasmodium parasitism predict juvenile survival?

#### Does *Plasmodium* parasitism predict juvenile return?

We investigated juvenile return separately by sex (males = 73, females = 72), using the following predictors: *Plasmodium* infection status, parasite load, body condition, year, wing length, and the interaction between body condition and parasite load as a potential predictor. No models outcompeted the null model for juvenile female return; see Table J in the S1 Supporting Tables. For juvenile males, one model was ranked higher than the null, although the null was included in the top model set <2 ΔAICc; see Table K in the S1 Supporting Tables. This top model showed that juvenile males with longer wings were more likely to return than were males with shorter wings (estimate = 0.22 ± 0.14 SE); for each mm increase in wing length, males had 1.24 times higher odds of return. See S1 Supporting Analyses Section B for an additional analysis predicting juvenile return from hatch date.

#### Does *Plasmodium* parasitism predict lifespan in returning juveniles?

Predictors of lifespan in returning juveniles (n = 72) included: wing length (centered by sex), centered body condition (calculated separately by sex), sex, infection status, parasite load, first year breeding, and all potential interactions. We retained five top models 2 ΔAICc; see Table L in the S1 Supporting Tables. Conditional model averaging showed that females have 1.78 times shorter lifespans than males (estimate =-0.57 ± 0.16 SE). Regardless of sex, each unit of increased body condition resulted in 1.14 times shorter lifespan (estimate =-0.13 ± 0.06 SE), and each mm increase in wing length was associated with 1.10 times shorter lifespan (estimate =-0.10 ± 0.05 SE). Birds with juvenile infections had 1.10 times shorter lifespans (estimate = - 0.23 ± 0.18 SE) than those that were uninfected as juveniles; each unit increase in *Plasmodium* load was associated with a 1.0005 times shorter lifespan (estimate =-5.3^-4^ ± 4.2^-4^ SE). See Figure 6 and Table L in the S1 Supporting Tables for additional details.

**Figure 6.** Model averaged coefficients from a model comparison identifying predictors of lifespan (in years) of juvenile juncos (*Junco hyemalis carolinensis*) that returned to the Mountain Lake Biological Station population (Pembroke, VA; n = 72). Predictors reflect data collected during the juvenile stage. Shown are estimates derived from conditional averaging of generalized linear models with a log link, accompanied by the unconditional standard error. ‘Infected’ refers to a diagnosis of *Plasmodium* infection, and ‘load’ refers to the relative *Plasmodium* load quantified by qPCR; see Methods and Materials for details.

## Discussion

Despite its popularity as a model system, the impacts of *Plasmodium* on avian host fitness remain unclear, especially in songbird populations that are historically sympatric with mosquitoes that vector *Plasmodium*. Here we investigated various metrics of host survival (return to the population and documented lifespan) and reproduction (fledging success and first-year breeding) in chronically infected adults and juveniles with acute infections. We hypothesized that negative impacts of *Plasmodium* infections on junco survival and reproduction would vary by host age and sex. Broadly, these hypotheses are supported, as juvenile males and first-year females likely experience the highest probability of *Plasmodium*-related mortality, while juvenile infections may reduce induce terminal investment in breeding in young adults.

### Plasmodium infection and load in adult juncos

The best predictor of *Plasmodium* infection in adult juncos was advanced age, aligning with results from previous studies in songbirds, including juncos [45–47,47–50]. Specifically, we found a higher probability of *Plasmodium* infection among third year juncos relative to yearlings. We also noted a higher probability of infection among females than males.

An important exception, however, is that we also detected lower probability of infection in second year females compared to yearling females. This could be explained by an increased probability of dispersal, mortality, or infection clearing in infected females during their second winter. We suggest that the most likely explanation is that the interaction of *Plasmodium* infection and first year breeding efforts leads to subsequent elevated overwinter mortality in this group. In a closely related songbird, the song sparrow (*Melospiza melodia*), female survivorship declined with increased breeding effort (i.e., days spent providing parental care) and the decline was greater in younger females [51] potentially because younger females had reduced access to high quality mates and food (ibid). Similarly, female blue tits showed a negative relationship between malaria parasite resistance and reproductive effort, but only in yearling females [52].

Therefore, it is possible that the added burden of *Plasmodium* infection during a female’s first breeding season may increase mortality in this demographic, leading to a relatively low probability of infection among second year females.

Variation in parasite load among infected adults was best explained by sampling year, more so than by sex or adult age category. Year-based variation in load likely reflects the influence of environmental factors, such as weather and food availability. These factors may also affect overwinter survival [53,54] and timing of onset of breeding [55]. This suggests that endocrine dynamics may influence yearly variation in load. For example, corticosterone, which increases *Plasmodium* parasite load [56], is highest in juncos during the early breeding season [57].

Therefore, unusually high loads such as those observed in 2006 may indicate a relatively early start to the breeding season, or perhaps some other phenomenon associated with increased corticosterone.

### Plasmodium infection and load in juvenile juncos

Environmental factors may also influence the probability of infection in juveniles, as year was a strong predictor of infection. The probability of *Plasmodium* infection was higher in 2003 and 2009, as compared to 2012, the year when it was lowest. It is possible that vector abundance was elevated during these years, although additional data are needed to test this hypothesis. The relatively small number of infected juveniles in our study reduced our ability to detect a relationship between juvenile parasite load and year; however, we would predict higher loads in years where food availability is lower, as nutrition supplementation has been shown to reduce parasitemia [34,58]. Future research with a larger sample size is needed to better understand which specific environmental factors explain variation in infection probability and *Plasmodium* load in juvenile songbirds.

Interestingly, we detected a female bias in *Plasmodium* infection status among juveniles, as was shown in adults. It is possible that young female birds are more attractive to vectors than young males; however, an experimental study using nestling great tits (*Parus major*) showed no host sex preference in feeding patterns of *Culex pipiens* mosquitoes [59], known vectors of *Plasmodium* [60]. Alternatively, mortality may have been higher in *Plasmodium*-infected nestling and juvenile males. The latter possibility is perhaps more probable, as male nestlings may show reduced immune function compared to females [29,61,62]. Similarly, males may show lower body condition when faced with haemosporidian [28] or ectoparasite [29] infections.

### Plasmodium parasitism and adult junco reproductive success

We predicted that *Plasmodium* infections in adult juncos would be associated with higher probability of nest failure. This prediction was not supported, as none of the top models predicting fledging success among adult social pairs included *Plasmodium*-related metrics. However, it remains possible that especially high loads or certain *Plasmodium* lineages might prevent adults from securing a mate altogether and this possibility should be addressed in future studies.

Interestingly, the best predictor of fledging success was male wing length. In juncos, male wing length increases notably from one to two years of age [63], and females tend to prefer older males as mates [64]. Thus, the relationship between successful fledging and male wing length could reflect higher success in older males. However, age was not selected as a predictor of fledging success in our model comparison. An alternative is that wing length could be related to territory quality or quality of male parental care, which includes nestling provisioning and defense against predators [65].

### Juvenile Plasmodium parasitism and first-year reproductive success

Among returning juveniles, we predicted that *Plasmodium* infection would be associated with reduced probability of first-year breeding. Surprisingly, we observed the opposite; infected juveniles with relatively higher *Plasmodium* loads and relatively heavier bodies showed an increased probability of first year breeding. One possibility is that individuals with heavy bodies were able to recover from high intensity infections and therefore had the energy and nutrient resources to invest in first year breeding. If this were true, however, we would predict body condition alone to be selected as an important predictor of first year breeding, but this was not the case. Rather, we suggest that terminal investment is a more likely explanation; the terminal investment hypothesis predicts elevated reproductive effort in individuals with a shortened life expectancy [66]. Indeed, we observed shorter lifespans in returning juncos diagnosed with *Plasmodium* infections as juveniles, compared to those that were uninfected. Furthermore, terminal investment has been demonstrated in a close relative of the junco, the song sparrow [67].

Our final prediction for this topic was that we would observe stronger *Plasmodium*-related decreases in reproductive success in juvenile juncos, relative to adults. It is unclear whether this hypothesis is supported; while we found no evidence of *Plasmodium*-related decreases in adult reproductive success, further research is needed to disentangle the effects of juvenile *Plasmodium* infections and body condition on lifetime reproductive success. However, female collared flycatchers (*Ficedula albicollis*) produced fewer recruited young in their lifetime if they were infected with either *Plasmodium* or *Haemoproteus* during their first year [33], which supports the possibility of similar effects in juncos.

### Plasmodium infection and adult junco survival

Among adults, we predicted that *Plasmodium* infection would predict a reduced probability of return (a proxy of survival), especially in females. This prediction was not supported, as neither infection status nor parasite load were a strong predictor of return in either sex. However, females paired with an infected male had a relatively higher probability of return, indicating that females may be indirectly impacted by male infections. We suggest that females with uninfected partners are less likely to survive the winter than females with infected mates. If uninfected males provide poor parental care, for example because they prioritize extrapair copulation, this could cause females to expend additional resources to compensate. Indeed, female juncos are known to compensate for reduced parental care from their mates [68–70]. Elevated reproductive effort is likely to increase mass loss [71] and immunosuppression in females [72], possibly even leading to an increase in parasite load [73,74]. Further research into the relationship between female return and male infection status is warranted.

### Plasmodium infection and juvenile junco survival

We predicted a reduced probability of return and shorter lifespans in *Plasmodium*-infected juveniles, especially males. Neither *Plasmodium* infection nor load alone was a significant predictor of return in juveniles of either sex. *Plasmodium* itself may not be an important predictor of juvenile mortality in this population, yet we cannot rule out the possibility that *Plasmodium*-related mortality occurred in young birds (e.g. nestlings and new fledglings) before we were able to sample them during juvenile census.

On the other hand, among returning juveniles, those that carried infections lived shorter lives than those without *Plasmodium* infection as juveniles; there was also a negative relationship between juvenile load and lifespan. This aligns with earlier work showing *Plasmodium*-related reductions in lifespans of great reed warblers (*Acrocephalus arundinaceus*) [7]. Similarly, Seychelles warblers contracting *Haemoproteus*, another malaria-causing parasite, also showed decreased survival compared to those without juvenile infections [75]. We also observed a higher probability of infection in juveniles that hatched earlier in the season, likely because earlier hatching birds had more opportunities for vector exposure. It is unclear whether this elevated risk of infection constitutes a cost of early hatching or whether it may be beneficial. For example, if *Plasmodium* risk is highest in the autumn, as has been suggested [76], it may benefit young birds to contract *Plasmodium* early enough in the season to recover from acute infection before winter. Further work is needed to address the interaction of hatch date and *Plasmodium* infection on juvenile songbird recruitment.

## Conclusion

Using a long-term dataset from a North American songbird population with a high prevalence of *Plasmodium*, we found evidence of both direct and indirect relationships between infection and host fitness. Our results suggest that *Plasmodium* is most likely to reduce fitness in younger birds and that these effects vary by sex. Specifically, juvenile males may be more likely to die from *Plasmodium* than juvenile females, and yearling females may be at increased risk of mortality after their first breeding season if they were infected as juveniles. This latter pattern, along with a shorter estimated lifespan for infected juveniles, supports the probability of terminal investment in juncos that were infected with *Plasmodium* as juveniles. Additional research is needed to understand sex differences in the impact of *Plasmodium* on juvenile survival.

For adults, we found that *Plasmodium* is unlikely to reduce survival or reproductive success directly. However, females may be indirectly impacted by male infection status, as adult females of any age mated to uninfected males showed reduced probability of returning to the population. Further research is needed to determine whether *Plasmodium* infection itself is associated with the quality of male parental care, or if susceptibility and/or response to *Plasmodium* infection is coincidentally linked to other traits that make certain males better breeding partners.

Here, we aimed to identify *Plasmodium* impacts on host fitness that have been understudied. We note, however, that our approach, as with most avian field studies, is subject to capture bias of relatively healthy birds [77] and may underestimate *Plasmodium* prevalence and the true impact of infection on songbird fitness. For example, it remains unclear what proportion of young juncos died from *Plasmodium* before they could be captured as juveniles, It is also possible that *Plasmodium* infections and/or relatively higher loads are more common in adults that fail to secure a mate. Other key limitations for our approach include possible underestimates of offspring number, as some individuals may have built nests outside of the study area, and possible false negatives in our *Plasmodium* diagnostics due to genetic variation among parasite lineages. In addition, our sample sizes were modest, especially for the infected juvenile demographic. Given these limitations, our results should be interpreted as conservative.

Nevertheless, we provide evidence of reduced songbird fitness associated with *Plasmodium* parasitism. Given the potential for virulent *Plasmodium* lineages to spread [78], and the potential for climate change to lengthen the active season [79], increase the abundance, and widen the distribution of *Plasmodium*-vectoring mosquitoes [80–82], continued research on this topic is warranted. In conclusion, despite an ancient coevolutionary relationship [2,83] our work suggests that *Plasmodium* may continue to have a selective impact on wild songbird populations to this day.

### Research ethics statement

Fieldwork was conducted under scientific collecting permits from the Virginia Department of Game and Inland Fisheries, and the US Fish and Wildlife, as well as USGS Federal Bird Banding Permit No. 20261. All methods were approved by the Indiana University Bloomington Institutional Animal Care and Use Committee.

## Acknowledgments

We thank Ketterson lab members, past and present, for their efforts in collecting junco data and samples at Mountain Lake Biological Station, as well as MLBS staff that facilitated this project. We also thank James Adelman and Arietta Fleming-Davies for consultation on statistical analyses. We also thank Daniel Becker, who co-advised CM during her time as an NSF REU trainee.

## Author Contributions

KMT conceived the ideas of this study and designed the methodology. EDK managed the long-term study yielding the data and samples for this project. KMT secured funding for laboratory analysis of blood samples. CM and KMT prepared molecular samples for analysis. OVG designed the molecular methodology and collected the molecular data. KMT and EDK analyzed the data and led the writing of the manuscript. All authors contributed critically to the drafts and gave final approval for publication.

## Data Availability Statement

Data and code associated with this study are available at https://zenodo.org/records/15376411

## Competing of Interests

The authors have no competing interests to declare.

## Supporting Information Captions

S1 Supporting Text. Supplementary information about the study system (Section A), junco capture and morphometric data collection (Section B), sample selection (Section C), *Plasmodium* diagnostics in junco blood (Section D), and junco sexing PCR (Section E).

S1 Supporting Tables. Results of AICc model comparisons investigating *Plasmodium* parasitism and other ecological factors as predictors of junco fitness metrics; details are provided for models within two ΔAIC. See code for all candidate models.

S1 Supporting Analyses. Additional model comparison analyses investigating hatch date as a predictor of *Plasmodium* infection in juvenile juncos (Section A) and investigating whether *Plasmodium* parasitism predicts juvenile return in juncos with known hatch dates (Section B).

**S1 Tables**. Results of AICc model comparisons investigating *Plasmodium* parasitism and other ecological factors as predictors of junco fitness metrics; details are provided for models within two ΔAICc. See code for all candidate models.

**Table A.** Generalized linear models predicting whether adult dark-eyed juncos (*Junco hyemalis carolinensis*) are infected with *Plasmodium*. All models use a binomial distribution. Shown are parameter estimates ± standard error. Age categories include individuals in their first breeding season (1 yr old), those two years of age, and those three years of age or older. See Methods and Materials for details on *Plasmodium* diagnostics. See code file for lists of all candidate models.

| Model | Intercept | Sex (F) | Age category (2 yr) | Age category (3 yr +) | Age category (2): Sex (F) | Age category (2): Sex (F) | k | <u>AICc</u> | <u>$\Delta$ AICc</u> | Weight |
| --- | --- | --- | --- | --- | --- | --- | --- | --- | --- | --- |
| 1.4 | -1.39 | | | | -1.76 ± 0.99 | 14.49 ± 1073.11 | 6 | 139.27 | 0.00 | 0.46 |
| 2.1 | -0.85 | 0.67 ± 0.45 | -0.10 ± 0.46 | 1.69 ± 0.62 | | | 4 | 140.49 | 1.22 | 0.25 |
| 1.3 | -0.43 | | -0.13 ± 0.46 | 1.41 ± 0.58 | | | 3 | 140.69 | 1.42 | 0.23 |
| Null | -0.19 |  |  |  |  |  | 1 | 145.25 | 5.98 | 0.02 |

**Table B.** Generalized linear models predicting parasite load in adult dark-eyed juncos (*Junco hyemalis carolinensis*) infected with *Plasmodium*. All models use a negative binomial distribution. Shown are parameter estimates ± standard error. Age categories include individuals in their first breeding season (1 yr old), those two years of age, and those three years of age or older. The reference year is 2000. Wing length is centered by sex. See Methods and Materials for details on *Plasmodium* diagnostics. See code file for lists of all candidate models.

| Model | Intercept | Sex (F) | Year (1999) | Year (2006) | Year (2009) | Year (1999): Sex (F) | Year (2006): Sex (F) | Year (2009): Sex (F) | Wing | k | AICc | $\Delta$ AICc | Weight |
| --- | --- | --- | --- | --- | --- | --- | --- | --- | --- | --- | --- | --- | --- |
| 1.4 | 2.62 | | 1.35 ± 0.55 | 2.20 ± 0.51 | 1.86 ± 0.49 | | | | | 5 | 467.19 | 0.00 | 0.27 |
| 2.5 | 2.10 | 0.69 ± 0.37 | 1.30 ± 0.54 | 2.64 ± 0.51 | 1.96 ± 0.49 | | | | | 6 | 467.35 | 0.16 | 0.25 |
| 1.1 | 1.92 | | | | | 0.89 ± 1.03 | -2.29 ± 0.97 | 0.05 ± 0.93 | | 9 | 467.50 | 0.31 | 0.23 |
| 2.4 | 2.61 | | 1.07 ± 0.55 | 2.36 ± 0.51 | 1.95 ± 0.49 | | | | -0.22 ± 0.12 | 6 | 467.51 | 0.32 | 0.23 |
| Null | 4.22 |  |  |  |  |  |  |  |  | 2 | 475.83 | 8.64 | 0.00 |

**Table C.** Generalized linear models predicting whether juvenile dark-eyed juncos (*Junco hyemalis carolinensis*) are infected with *Plasmodium*. All models use a binomial distribution. Shown are parameter estimates ± standard error. Body condition (‘body cond.’) was quantified as residuals of a linear regression of mass against tarsus length, with separate models for each sex. Reference year is 2012. See Methods and Materials for details on *Plasmodium* diagnostics. See code file for lists of all candidate models.

| Model | Intercept | Sex (F) | Year (2000) | Year (2003) | Year (2004) | Year (2006) | Year (2007) | Year (2009) | Year (2010) | Year (2014) | Body cond. | k | AICc | $\Delta$ AICc | Weight |
| --- | --- | --- | --- | --- | --- | --- | --- | --- | --- | --- | --- | --- | --- | --- | --- |
| 6 | -3.39 | 0.72 ± 0.45 | 1.31 ± 1.29 | 3.34 ± 1.16 | 2.07 ± 1.19 | 1.92 ± 1.13 | 0.37 ± 1.46 | 4.22 ± 1.25 | 1.65 ± 1.22 | 1.21 ± 1.29 | | 10 | 155.78 | 0.00 | 0.45 |
| 4 | -2.89 | | 1.19 ± 1.28 | 3.14 ± 1.14 | 1.88 ± 1.18 | 1.75 ± 1.12 | 0.33 ± 1.46 | 3.99 ± 1.22 | 1.59 ± 1.22 | 1.19 ± 1.28 | | 9 | 156.12 | 0.34 | 0.38 |
| 8 | -3.39 | 0.72 ± 0.45 | 1.25 ± 1.30 | 3.42 ± 1.17 | 2.01 ± 1.20 | 1.97 ± 1.13 | 0.29 ± 1.47 | 4.15 ± 1.25 | 1.64 ± 1.22 | 1.18 ± 1.29 | 0.15 ± 0.22 | 11 | 157.65 | 1.87 | 0.18 |
| Null | -1.04 |  |  |  |  |  |  |  |  |  |  | 1 | 168.84 | 13.06 | 0.00 |

**Table D.** Generalized linear models predicting parasite load in juvenile dark-eyed juncos (*Junco hyemalis carolinensis*) infected with *Plasmodium* (n = 38). All models use a negative binomial distribution. Shown are parameter estimates ± standard error. Body condition (‘body cond.’) was quantified as residuals of a linear regression of mass against tarsus length, with separate models for each sex. Reference year is 2009. See Methods and Materials for details on *Plasmodium* diagnostics. See code file for lists of all candidate models.

| Model | Intercept | Sex (F) | Year (2000) | Year (2003) | Year (2004) | Year (2006) | Year (2007) | Year (2010) | Year (2012) | Year (2014) | Body cond. | k | <u>AICc</u> | <u>$\Delta$ AICc</u> | Weight |
| --- | --- | --- | --- | --- | --- | --- | --- | --- | --- | --- | --- | --- | --- | --- | --- |
| Null | 5.76 |  |  |  |  |  |  |  |  |  |  | 2 | 512.46 | 0.00 | 0.34 |
| 2 | 5.74 | | | | | | | | | | -0.25 ± 0.22 | 3 | 513.51 | 1.05 | 0.20 |
| 4 | 4.75 | | -1.26 ± 0.78 | -0.23 ± 0.47 | -3.71 ± 0.62 | -0.15 ± 0.50 | 0.72 ± 1.05 | 0.32 ± 0.66 | -1.52 ± 1.05 | -1.80 ± 0.78 | | 10 | 513.53 | 1.06 | 0.20 |
| 3 | 5.90 | -0.27 ± 0.40 | | | | | | | | | | 3 | 514.39 | 1.93 | 0.13 |

**Table E.** Generalized linear models predicting whether adult dark-eyed junco (*Junco hyemalis carolinensis*) social pairs (n = 52) raise at least one offspring to independence. All models use a binomial distribution. Shown are parameter estimates ± standard error. Male body condition was quantified as residuals of a linear regression of mass against tarsus length, and wing length was measured in mm. See Methods and Materials for details on *Plasmodium* diagnostics. See code file for lists of all candidate models.

| Model | Intercept | Male wing length | Male parasite load | Male body condition | k | AICc | $\Delta$ AICc | Weight |
| --- | --- | --- | --- | --- | --- | --- | --- | --- |
| 1.3 | -29.61 | $0.35 \pm 0.16$ | | | 2 | 70.78 | 0.00 | 0.21 |
| 3.8 | -40.26 | $0.48 \pm 0.19$ | $1.6^{-3} \pm 3.8^{-3}$ | $-0.68 \pm 0.35$ | 4 | 70.93 | 0.15 | 0.19 |
| Null | -0.15 |  |  |  | 1 | 73.86 | 3.08 | 0.04 |

**Table F.** Generalized linear models the number of fledged offspring produced by adult junco (*Junco hyemalis carolinensis*) social pairs that have raised at least one offspring (n = 24). All models use a Poisson distribution. Shown are parameter estimates ± standard error. See Methods and Materials for details on *Plasmodium* diagnostics. See code file for lists of all candidate models.

| Model | Intercept | Male parasite load | k | AICc | $\Delta$ AICc | Weight |
| --- | --- | --- | --- | --- | --- | --- |
| Null | 1.22 |  | 1 | 86.01 | 0.00 | 0.27 |
| 1.8 | 1.18 | $1.3^{-3} \pm 1.8^{-3}$ | 2 | 87.87 | 1.86 | 0.11 |

**Table G.** Generalized linear models predicting whether juvenile dark-eyed juncos (*Junco hyemalis carolinensis*; n = 72) that returned to the MLBS population attempted to breed in their first adult year. Predictive data were collected during each junco’s hatch year, during the annual juvenile census (see Methods and Materials for details). All models use a binomial distribution. Body condition was quantified as residuals of a linear regression of mass against tarsus length, with separate models run for each sex. See Methods and Materials for details on *Plasmodium* diagnostics. See code file for lists of all candidate models.

| Model | Intercept | Body condition *<br>parasite load | k | AICc | Δ AICc | Weight |
| --- | --- | --- | --- | --- | --- | --- |
| 11 | -1.27 | 0.02 ± 0.01 | 4 | 78.31 | 0.00 | 0.75 |
| Null | -1.03 |  | 1 | 85.16 | 6.85 | 0.02 |

**Table H.** Generalized linear models predicting whether adult female dark-eyed junco (*Junco hyemalis carolinensis*; n = 52) returned to the MLBS population. Return is used as a conservative proxy of survival. All models use a binomial distribution. Shown are parameter estimates ± standard error. See Methods and Materials for details on *Plasmodium* diagnostics. See code file for lists of all candidate models.

| Model | Intercept | Mate infection status (infected) | Parasite load | Mate age category (2 yrs) | Mate age category (3+ yrs) | k | AICc | $\Delta$ AICc | Weight |
| --- | --- | --- | --- | --- | --- | --- | --- | --- | --- |
| 1.5 | -1.61 | 1.61 ± 0.65 | | | | 2 | 61.78 | 0.00 | 0.24 |
| 2.5 | -1.45 | 1.65 ± 0.67 | 9.4<sup>-3</sup> ± 8.7<sup>-3</sup> | | | 3 | 62.13 | 0.36 | 0.20 |
| 1.7 | -2.64 | | | 2.02 ± 1.14 | 2.52 ± 1.14 | 3 | 63.25 | 1.48 | 0.11 |
| Null | -0.81 |  |  |  |  | 1 | 66.27 | 4.50 | 0.03 |

**Table I.** Generalized linear models predicting whether adult male dark-eyed junco (*Junco hyemalis carolinensis*; n = 52) returned to the MLBS population. Return is used as a conservative proxy of survival. All models use a binomial distribution. Shown are parameter estimates ± standard error. See Methods and Materials for details on *Plasmodium* diagnostics. See code file for lists of all candidate models.

| Model | Intercept | Fledging success<br>(successful) | Parasite load | Infection status<br>(infected) | k | AICc | $\Delta$ AICc | Weight |
| --- | --- | --- | --- | --- | --- | --- | --- | --- |
| 1.2 | -0.75 | $1.26 \pm 0.58$ | | | 2 | 71.16 | 0.00 | 0.22 |
| 2.4 | -0.89 | $1.31 \pm 0.59$ | $3.91^{-3} \pm 3.82^{-3}$ | | 3 | 72.31 | 1.15 | 0.12 |
| 2.1 | -0.56 | $1.31 \pm 0.59$ | | $-0.50 \pm 0.60$ | 3 | 72.72 | 1.55 | 0.10 |
| Null | -0.15 |  |  |  | 1 | 73.86 | 2.69 | 0.06 |

**Table J.** Generalized linear models predicting whether juvenile female dark-eyed junco (*Junco hyemalis carolinensis*; n = 72) returned to the MLBS population. Return is used as a conservative proxy of survival. All models use a binomial distribution. Shown are parameter estimates ± standard error. See Methods and Materials for details on *Plasmodium* diagnostics. See code file for lists of all candidate models.

| Model | Intercept | Infection status<br>(infected) | Parasite load | k | AICc | $\Delta$ AICc | Weight |
| --- | --- | --- | --- | --- | --- | --- | --- |
| Null | -0.06 |  |  | 1 | 101.81 | 0.00 | 0.19 |
| 3 | -0.28 | $0.76 \pm 0.53$ | | 2 | 101.82 | 0.01 | 0.19 |
| 5 | -0.11 | | $6.54^{-4} \pm 1.13^{-3}$ | 2 | 103.59 | 1.77 | 0.08 |

**Table K.** Generalized linear models predicting whether juvenile male dark-eyed junco (*Junco hyemalis carolinensis*; n = 73) returned to the MLBS population. Return is used as a conservative proxy of survival. All models use a binomial distribution. Shown are parameter estimates ± standard error. Body condition was quantified as residuals of a linear regression of mass against tarsus length, and wing length was quantified in mm. See Methods and Materials for details on *Plasmodium* diagnostics. See code file for lists of all candidate models.

| Model | Intercept | Wing length | Parasite load | Body condition | Infection status (infected) | k | <u>AICc</u> | <u>$\Delta$ AICc</u> | Weight |
| --- | --- | --- | --- | --- | --- | --- | --- | --- | --- |
| 2 | -17.83 | $0.22 \pm 0.14$ | | | | 2 | 102.77 | 0.00 | 0.23 |
| Null | 0.03 |  |  |  |  | 1 | 103.24 | 0.47 | 0.18 |
| 10 | -17.43 | $0.21 \pm 0.14$ | $-8.64^{-4} \pm 1.12^{-3}$ | | | 3 | 104.32 | 1.55 | 0.10 |
| 6 | 0.10 | | $-9.16^{-4} \pm 1.10^{-3}$ | | | 2 | 104.63 | 1.86 | 0.09 |
| 3 | 0.03 | | | $0.16 \pm 0.20$ | | 2 | 104.66 | 1.89 | 0.09 |
| 8 | -18.30 | $0.23 \pm 0.14$ | | | $-0.28 \pm 0.57$ | 3 | 104.70 | 1.93 | 0.09 |

**Table L.** Generalized linear models lifespan (in years) of juvenile dark-eyed juncos (*Junco hyemalis carolinensis*) that returned to the MLBS population (n = 72). All models use a Poisson distribution. Shown are parameter estimates ± standard error. Body condition was quantified as residuals of a linear regression of mass against tarsus length, with separate models by sex. Wing length was measured in mm. See Methods and Materials for details on *Plasmodium* diagnostics. See code file for lists of all candidate models.

| Model | Intercept | Parasite load | Sex (F) | Body condition | Infection status (infected) | Wing length | k | AICc | $\Delta$ AICc | Weight |
| --- | --- | --- | --- | --- | --- | --- | --- | --- | --- | --- |
| 26 | 1.18 | $-5.22^{-4} \pm 4.19^{-4}$ | $-0.57 \pm 0.16$ | $-0.13 \pm 0.06$ | | | 4 | 255.89 | 0.00 | 0.22 |
| 24 | 1.20 | | $-0.56 \pm 0.16$ | $-0.12 \pm 0.06$ | $-0.22 \pm 0.18$ | | 4 | 256.17 | 0.28 | 0.19 |
| 25 | 1.22 | | $-0.57 \pm 0.16$ | | $-0.25 \pm 0.18$ | $-0.09 \pm 0.05$ | 4 | 256.64 | 0.75 | 0.15 |
| 27 | 1.20 | $-5.43^{-4} \pm 4.28^{-4}$ | $-0.59 \pm 0.16$ | | | $-0.10 \pm 0.05$ | 4 | 256.76 | 0.87 | 0.14 |
| 4 | 1.15 | | $-0.58 \pm 0.16$ | | | | 2 | 257.71 | 1.81 | 0.09 |
| 16 | 1.20 | | $-0.54 \pm 0.16$ | | $-0.26 \pm 0.18$ | | 3 | 257.79 | 1.89 | 0.09 |
| Null | 0.91 |  |  |  |  |  | 1 | 269.85 | 13.95 | 0.00 |

**S1 Analyses**. Additional model comparison analyses investigating hatch date as a predictor of *Plasmodium* infection in juvenile juncos (Section A) and investigating whether *Plasmodium* parasitism predicts juvenile return in juncos with known hatch dates (Section B).

A) *Which factors predict* Plasmodium *infection status in juveniles with known hatch dates?*

We predicted that hatch date could influence the probability of juveniles contracting *Plasmodium*, with those hatching earlier in the season having a higher probability of infection. Because hatch date data were not available for all individuals, we used a subset of juveniles with known hatch dates (n=75) for this analysis. Predictors included sex, hatch date (given as days since January 1^st^), and body condition (calculated separately by sex). Year was not included as a predictor in the second analysis because of the relatively small number of observations per year in the reduced dataset. Model selection returned three models <2 ΔAICc. The univariate model showing a negative relationship between hatch date and probability of infection had the strongest weight and was selected as the top model (estimate =-0.05 ± 0.02 SE). The odds of infection were 1.05 times lower for each day later in the season in which a juvenile hatched; see Figure S1. Increased probability of return in juveniles that hatched earlier in the season may be attributable to food availability. For example, juvenile great tits (*Parus major*) in the Netherlands are more likely to return when they hatch near the early-season peak in caterpillar abundance (Visser et al., 1998).

**Figure S1.**
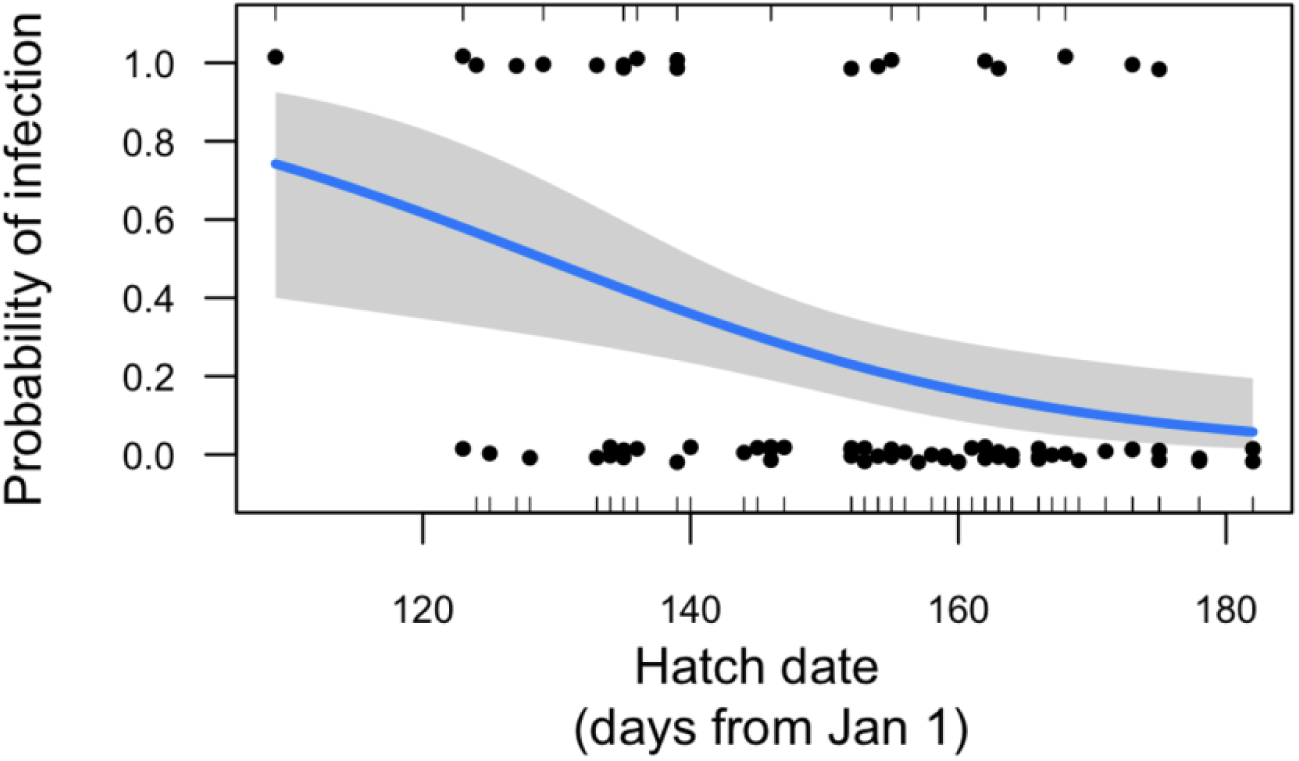
Juvenile dark-eyed juncos (*Junco hyemalis*; n=75) that hatch earlier in the breeding season near Mountain Lake Biological Station (Pembroke, VA) had a higher probability of *Plasmodium* infection than those that hatch later. Shown are estimates, accompanied by a 95% confidence interval, extracted from a generalized linear model with a binomial distribution. See Methods and Materials for *Plasmodium* diagnostic details. Hatch date is given as number of days since January 1^st^; for reference, May 1^st^ is Day 121 and July 1^st^ is Day 182. Points represent data for individual juncos and are jittered for easier viewing.

B) *Supplemental analysis: Does* Plasmodium *parasitism predict return in juveniles with known hatch dates?*

We added a third analysis to investigate whether *Plasmodium* parasitism predicted juvenile return in birds with known hatch dates (n=75). Predictors for this analysis included centered body condition (calculated separately by sex), wing length (centered by sex), sex, infection status, parasite load, and hatch date (given as number of days since January 1). Model selection returned three top models <2 ΔAICc explaining variation in juvenile return; each showed that individuals hatching earlier in the year were more likely to return to the population than those hatching later in the year, while two models also included *Plasmodium* diagnostics. Conditional model averaging revealed a 1.04 times lower probability of return in juveniles for each additional day in hatch date (estimate =-0.04 ± 0.02 SE), and a 2.74 times higher probability of return in *Plasmodium* infected juveniles compared to those that were uninfected (estimate = 1.01 ± 0.62 SE); see Figure S2. Probability of return was also higher with higher *Plasmodium* loads (estimate = 1.0 x 10^-3^ ± 0.001); however, the effect size was small, and the estimate interval crossed zero. The possibility of a non-linear effect of load on juvenile return cannot be ruled out.

**Figure S2.**
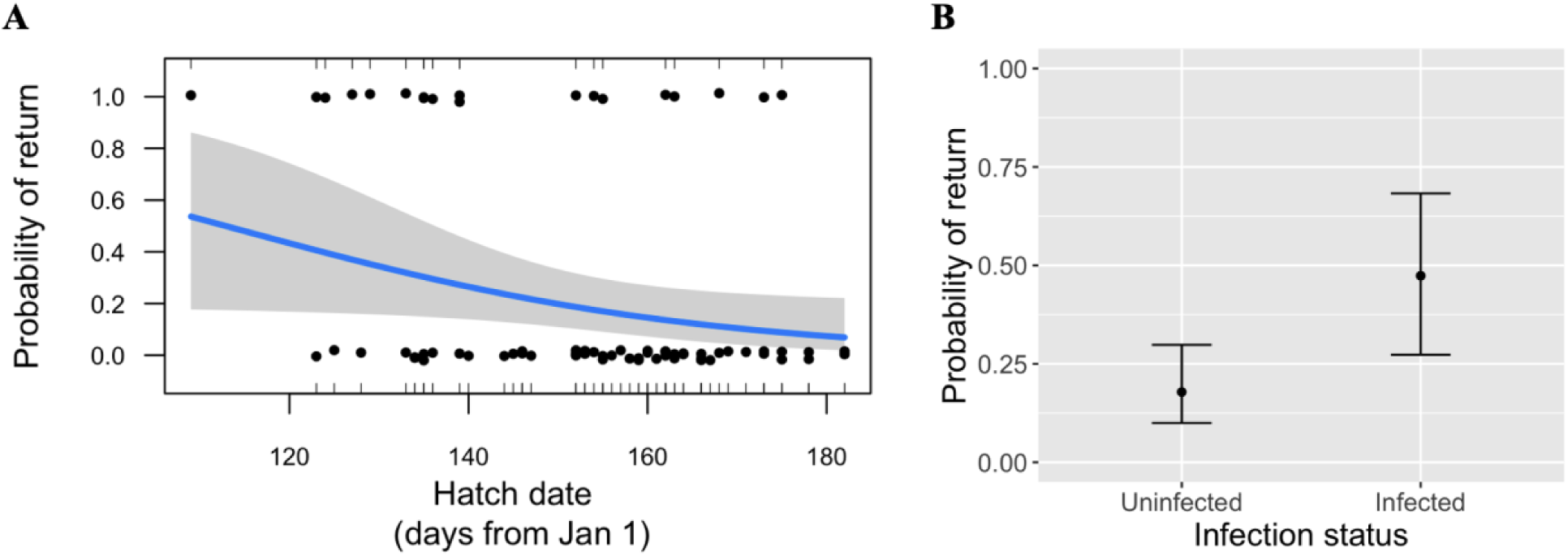
Juvenile dark-eyed juncos (*Junco hyemalis*; n=75) that hatch earlier in the breeding season near Mountain Lake Biological Station (Pembroke, VA) have a higher probability of return to the MLBS population than those that hatch later. Similarly, those that are infected with *Plasmodium* have a relatively higher probability of return. A) Shown are estimates with a 95% CI from a generalized linear model (binomial distribution) predicting probability of juvenile return from juvenile hatch date. Points represent data for individual juncos and are jittered for easier viewing. B) The proportion of returning juveniles is given for those with and without *Plasmodium* infection; each proportion is accompanied by a 95% CI. See Methods and Materials for *Plasmodium* diagnostic details. Hatch date is given as the number of days since January 1^st^; for reference, May 1^st^ is Day 121 and July 1^st^ is Day 182.

**S1 Supporting Text**. Supplementary information about the study system (Section A), junco capture and morphometric data collection (Section B), sample selection (Section C), *Plasmodium* diagnostics in junco blood (Section D), and junco sexing PCR (Section E).

A. *Study system*

<u>Host ecology</u> – Pair formation begins in late March-April, followed by nest building and egg laying (typically four eggs per nest) in April-early May (Nolan et al., 2002). Nests are built on or near the ground, and nest failure, often attributable to predation, is common (Ketterson et al., 1996). Breeding pairs renest following nest failure or successful fledging (Nolan et al., 2002).

Juncos are socially monogamous, with females providing incubation and both parents feeding nestlings (Nolan et al., 2002). After eggs hatch, young leave the nest after 11 – 12 days (Nolan et al., 2002). Fledglings are attended by parents for approximately two weeks, during which time they are provided additional food and warned of nearby predators. Following this period, offspring are considered independent of their parents (Wolf et al., 1988) and are termed ‘juveniles’. Juncos are relatively short-lived, with approximately 10% of fledged young, 42% of adult females, and 48% of adult males detected in the population in subsequent years (Ketterson et al., 1996; Wolf et al., 1988). Detecting an individual in subsequent years is termed ‘return’ and is a conservative proxy for survival.

<u>Parasite ecology</u> – All three haemosporidian genera, including at least four species of *Plasmodium* (*P. circumflexum*, *P. lutzi*, *P. elongatum*, and *P. cathemerium*) have been documented in the MLBS population (Slowinski et al., 2018). *Plasmodium* prevalence is high, with approximately 50 – 90% of adults diagnosed as infected via PCR, depending on the season (Becker et al., 2019; Slowinski et al., 2018, Talbott pers. obs.). *Plasmodium*-vectoring *Culex* mosquitoes are likely absent or at low density during early spring at this study site (Jackson & Paulson, 2006). Thus, adults sampled during early breeding season and diagnosed as *Plasmodium* positive probably acquired infections during the prior breeding season (or earlier) and were most likely in the chronic phase of infection when sampled. Infected juveniles, which were sampled later in the season, may have been carrying acute or chronic infections, depending on the timing of parasite exposure. It should be noted that *Plasmodium* can exit the host bloodstream and enter other tissues (Atkinson et al., 2008), causing false negatives in PCR diagnostics of host blood. Thus, our diagnoses should be considered conservative.

B. *Field methods and morphometric data collection*

During each census, birds were captured using baited mist-nets and walk-in traps. At first capture, each bird was given a unique combination of color bands and a United States Fish and Wildlife Service band for identification. Small blood samples were collected by brachial venipuncture (<2% blood volume estimated by bird mass). Blood samples were preserved with Longmire’s solution (Longmire et al., 1997). DNA extracted from preserved blood was used for *Plasmodium* diagnostics (see below) and for sexing non-returning juveniles (see Section E, below). For all captured birds, morphological data were recorded, including: mass measured with a Pesola spring scale (to nearest 0.1 g), flattened wing length measured with a ruler (to nearest mm), and tarsus length measured with calipers (to nearest 0.1 mm). Body condition was quantified as residuals of a linear regression of mass against tarsus length (Labocha & Hayes, 2012), with separate models by age class (juvenile or adult) and sex. During the adult census, some females were gravid, which skewed the mean of female mass; thus, we included only male body condition data in our analyses of adults.

C. *Sample selection*

For adults, we focused on samples and data collected from individuals during the adult census in the years 1999, 2000, 2006, and 2009. We selected data from years in which samples from 12 or more adult pairs were available. Because this population of juncos has been the subject of studies investigating the impact of testosterone implants on reproductive behavior (Ketterson et al., 1996), we only included samples from individuals that were not treated with testosterone during the year of sampling. To control for ecological conditions that varied by year (e.g., predator density, food availability, and weather), we selected approximately the same number of successful (i.e., raised at least one young to fledging) and unsuccessful pairs (no known fledged young and at least two failed nests) for each year. Only adult pairs that met one of these sets of criteria (‘successful’ or ‘unsuccessful’) were included.

For juveniles, we focused on data collected during the juvenile census. We selected samples from nine years between 2000 to 2014, in which nine or more juvenile samples were available per year. For juveniles with known parents, we included only one juvenile per brood. We also included only individuals with parents that were not treated with testosterone in the year the juvenile hatched. We balanced yearly samples by sex and return status (i.e., whether an individual was detected in the MLBS population in years following its hatch year) when possible. Sex for returners was confirmed by adult plumage, breeding condition (cloacal protuberance in males and brood patch in females), or breeding behavior (territorial singing in males and nest incubation in females). Sex in non-returners was confirmed using sexing PCR (see Section E for details). Some juveniles captured at census had already been banded as nestlings and therefore their hatch date was known (n=75); this subset of individuals was used for analyses in which hatch date was an independent variable (see Supplemental Analyses A and B).

D. Plasmodium *Diagnostics*

We used quantitative polymerase chain reaction (qPCR) to detect *Plasmodium* infections in junco blood samples and to assess relative amounts of parasite DNA and host DNA. We used primer set L9/NewR, which amplifies a 188-bp fragment of the *Plasmodium* cytochrome b gene (Knowles et al., 2010). To control for host DNA, we used the primer set SFSR3fb/SFSR3rb, which targets a 114-bp fragment of a single-copy nuclear gene that is conserved among vertebrate taxa (Asghar et al., 2011). DNA was quantified using a NanoDrop 1000 spectrophotometer (Thermo Scientific, MA, USA) and diluted to a working concentration of 3-8 ng/ul. The qPCR relative quantification experiment was done on Applied Biosystems® QuantStudio 3 Real-Time PCR System (Life Technologies, CA, USA). The qPCR reaction was conducted in volume 20 µl containing: 10 µl of GoTaq® qPCR Master Mix (Promega, WI, USA), and 300 nM of each (forward and reverse) primer.

Five microliters of template, standard (pooled DNA from juncos known to be *Plasmodium* infected) or water (for negative control) were added to the appropriate qPCR mix. Amplification was performed using the following cycles: 2 min at 95°C; 40 cycles of 15 s at 95°C, 30 s at 56°C and 72°C for 40 s. A dissociation curve was established following each qPCR reaction. Each reaction was run in triplicate and *Plasmodium* relative quantification was estimated by calculating the mean value for the triplicate. All dissociation curves were examined for the presence of nonspecific amplification or primer dimer formation. The standard sample containing both host and *Plasmodium* DNA was amplified with both primers on each plate for data normalization purposes. To calculate relative parasite load, we used a relative quantification metric reflecting the proportion of parasite DNA relative to a control sample.

We did not correct for differences in primer efficiency because both primer pairs had an efficiency of approximately 96% when DNA concentration ranged from 2.5 to 40 ng per reaction. We selected the sample with the lowest cycle threshold value for the gene of interest (i.e., the highest amount of parasite DNA) as the control. Relative quantification values are therefore bounded by 0 (an uninfected bird) to 1 (a bird with the same amount of parasite DNA relative to host DNA as the control). We assigned a load of zero to samples with cycle threshold values ≥ 37 for parasite amplification.

E. *Sexing PCR*

To confirm sex in non-returning juveniles, we ran PCR in 10 μL reactions. Each reaction included 1 μL undiluted DNA, 1 μL each of primers P2 and P8 at 10 μM, 2 μL nuclease-free water, and 5 μL Promega PCR Master Mix. The conditions included a denaturing step of 1 minute at 94°C, followed by 30 cycles of 94°C for 60 sec, 48°C for 60 sec, and 72°C for 2 min, followed by an extension phase at 72°C for 10 minutes (Port & Greeney, 2012). PCR products were visualized after running electrophoresis at 90 v for 1.5 hrs on a 3% agarose gel stained with GelRed (Biotium).

## Notes

https://zenodo.org/records/15376411

### Competing Interest Statement

The authors have declared no competing interest.

### Funding

- American Museum of Natural History: Chapman Research Grant
- National Institutes of Health: Common Themes in Reproductive Diversity T32HD049336
- Black Hills State University‘s Western SD Genetics and Genomics Core Facility
- U.S. National Science Foundation: 2050311

