## Supplementary figures and images for "Fitness impacts of *Plasmodium* vary by host age and sex in a North American songbird"

### Figure 1

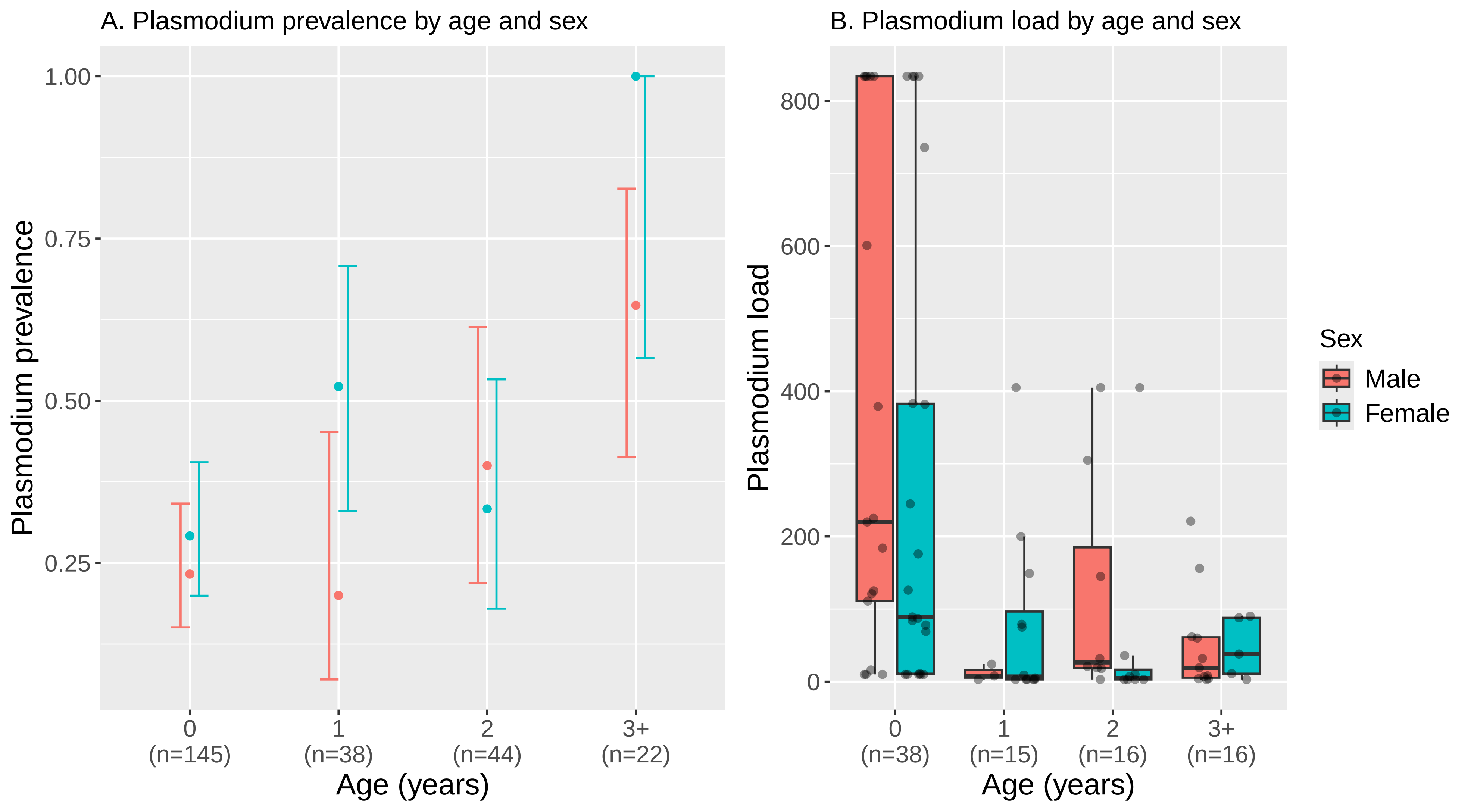

### Figure 2

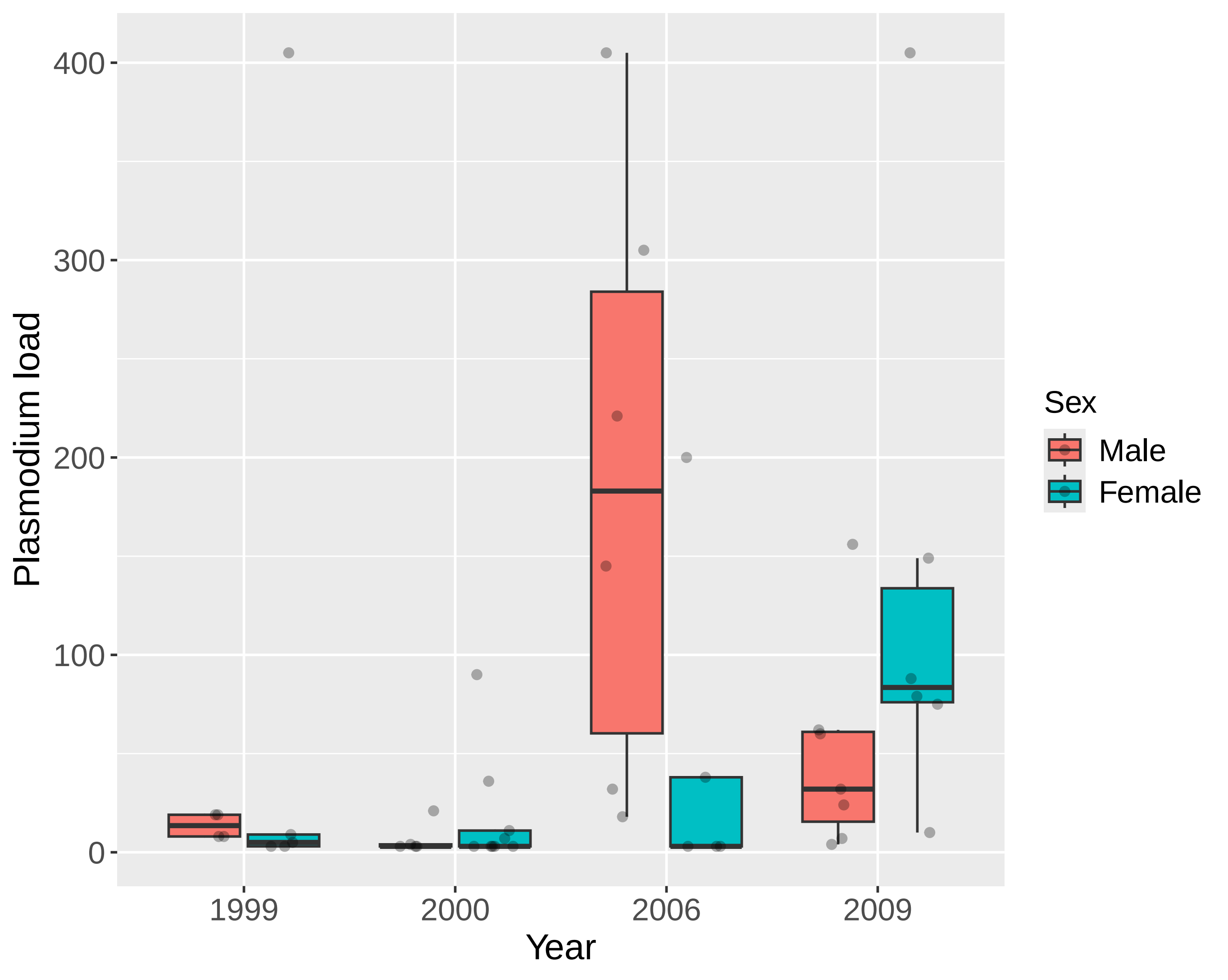

### Figure 3

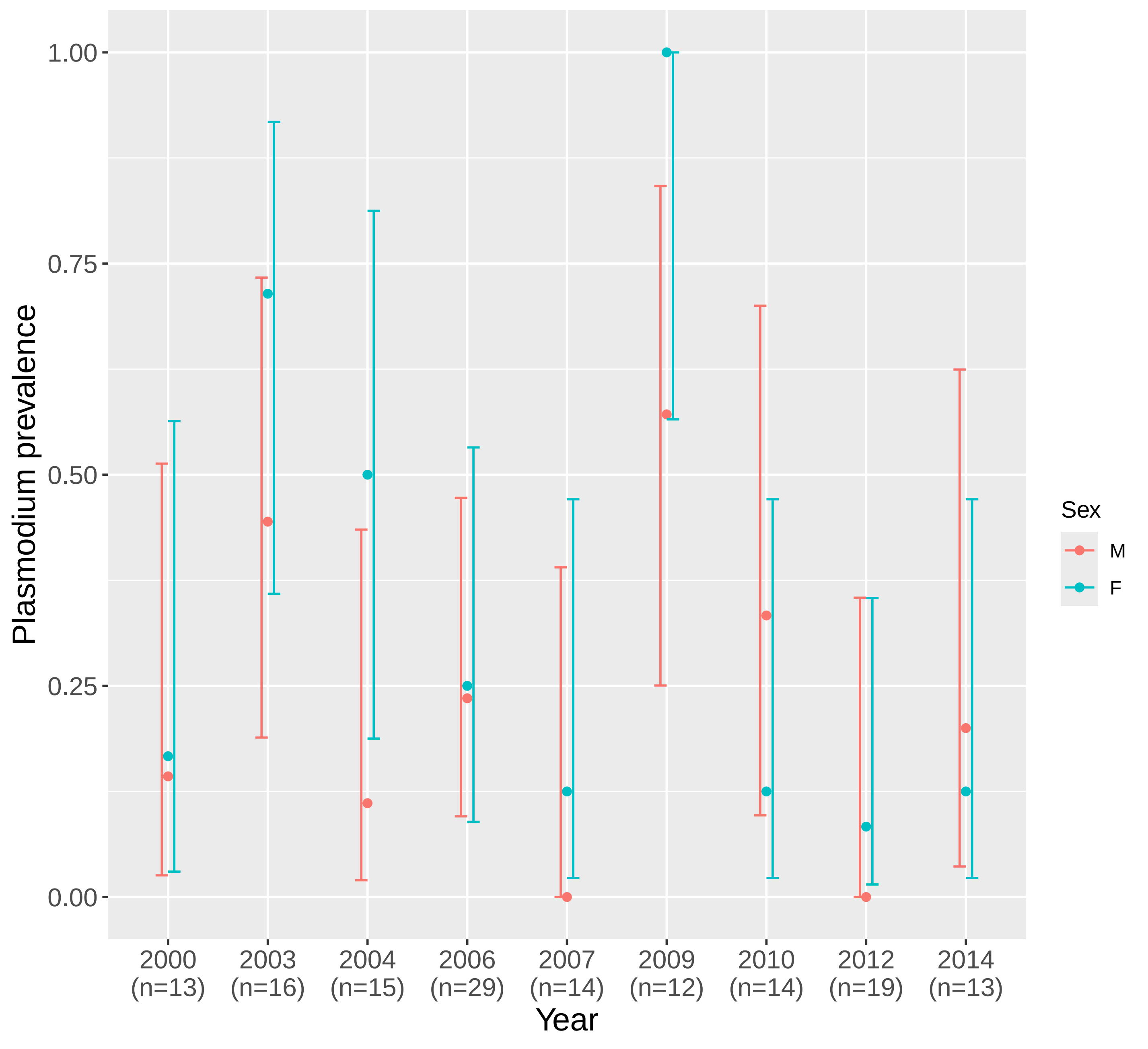

### Figure 4

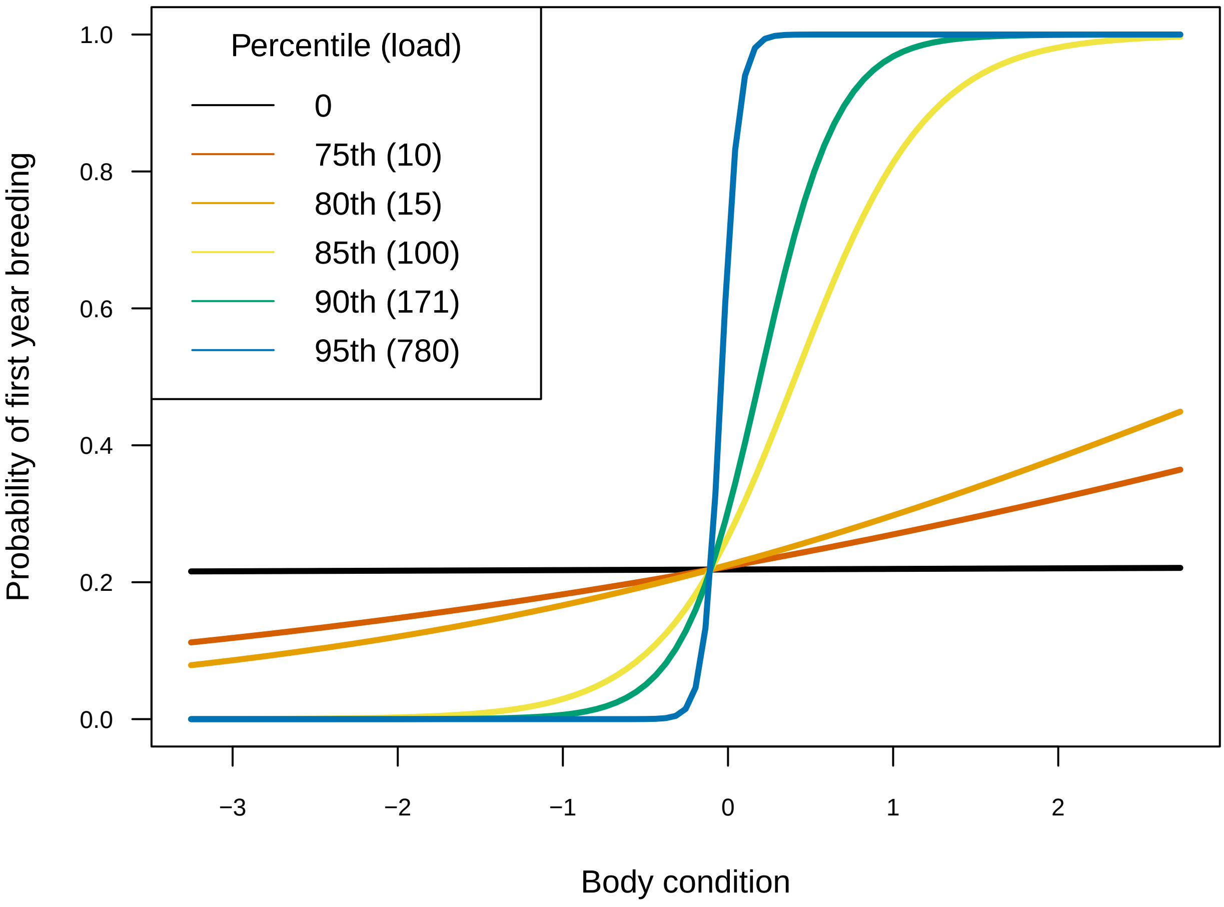

### Figure 5

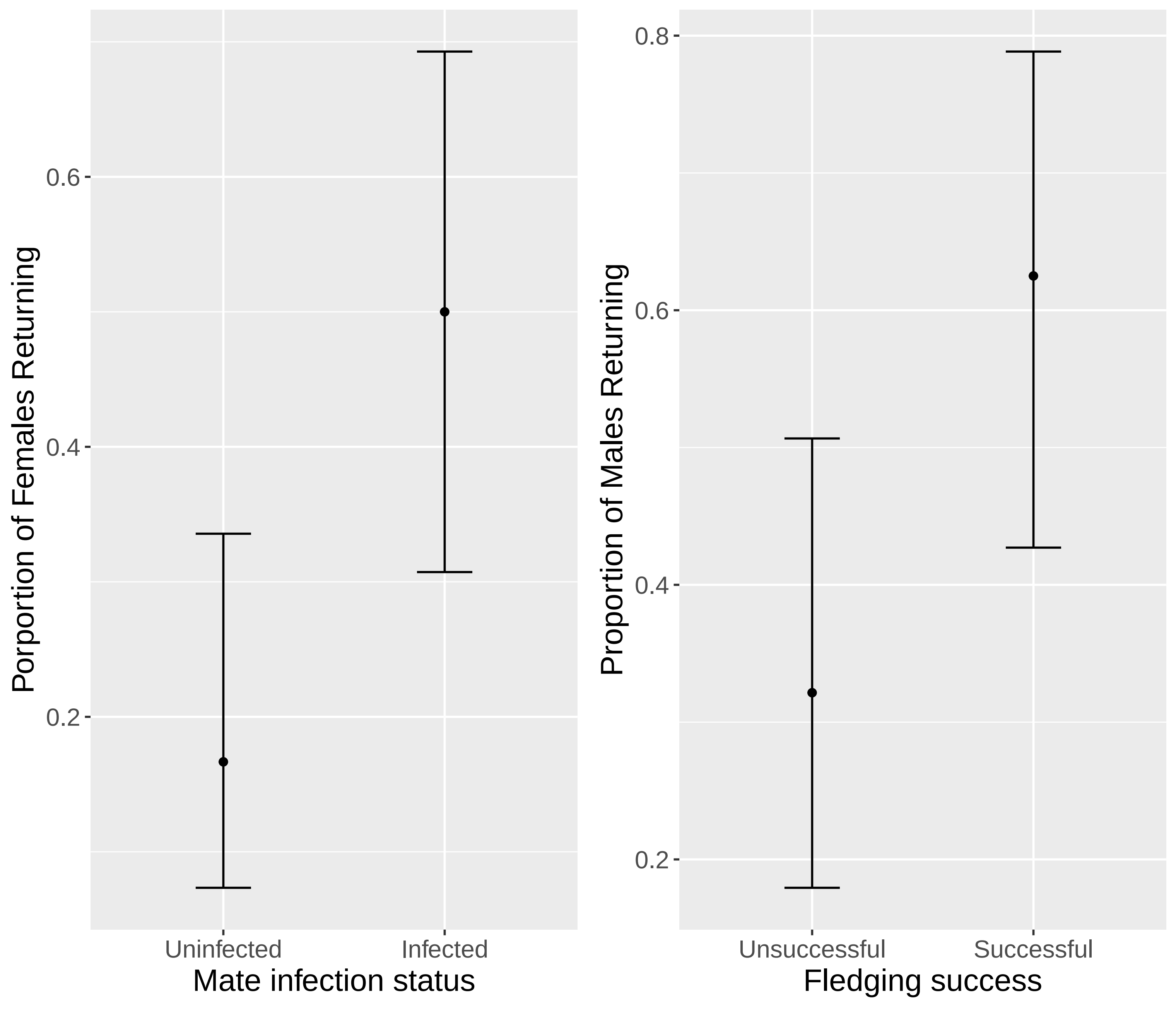

### Figure 6

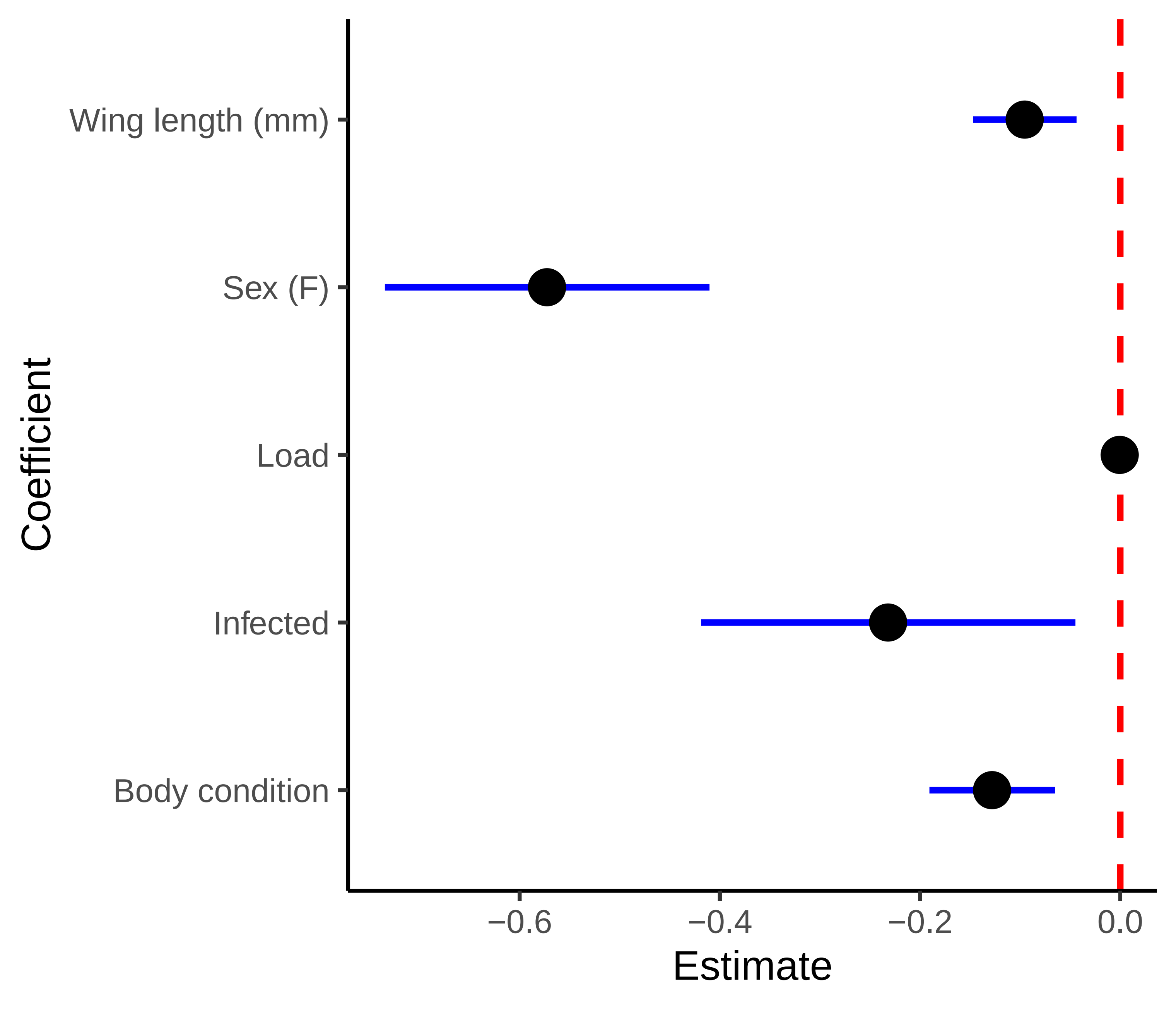
